# Salt-inducible kinase inhibition promotes the adipocyte thermogenic program and adipose tissue browning

**DOI:** 10.1101/2022.10.27.514130

**Authors:** Fubiao Shi, Flaviane de Fatima Silva, Dianxin Liu, Hari U Patel, Jonathan Xu, Wei Zhang, Clara Türk, Marcus Krüger, Sheila Collins

## Abstract

Graphical abstracts

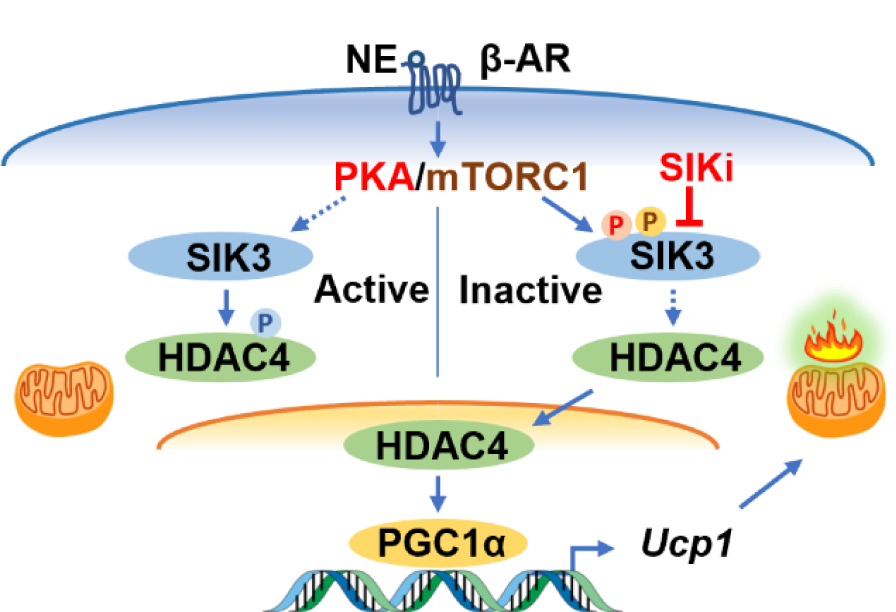

**Objective:** Norepinephrine stimulates the adipose tissue thermogenic program through a β-adrenergic receptor (βAR) – cyclic adenosine monophosphate (cAMP) – protein kinase A (PKA) signaling cascade. We discovered that a noncanonical activation of the mechanistic target of rapamycin complex 1 (mTORC1) by PKA is required for the βAR-stimulation of adipose tissue browning. However, the downstream events triggered by PKA-phosphorylated mTORC1 activation that drive this thermogenic response are not well understood.

**Methods:** We used a proteomic approach of Stable Isotope Labeling by/with Amino acids in Cell culture (SILAC) to characterize the global protein phosphorylation profile in brown adipocytes treated with the βAR agonist. We identified salt-inducible kinase 3 (SIK3) as a candidate mTORC1 substrate and further tested the effect of SIK3 deficiency or SIK inhibition on the thermogenic gene expression program in brown adipocytes and in mouse adipose tissue.

**Results:** SIK3 interacts with RAPTOR, the defining component of the mTORC1 complex, and is phosphorylated at Ser^884^ in a rapamycin-sensitive manner. Pharmacological SIK inhibition by a pan-SIK inhibitor (HG-9-91-01) in brown adipocytes increases basal *Ucp1* gene expression and restores its expression upon blockade of either mTORC1 or PKA. Short-hairpin RNA (shRNA) knockdown of *Sik3* augments, while overexpression of SIK3 suppresses, *Ucp1* gene expression in brown adipocytes. The regulatory PKA phosphorylation domain of SIK3 is essential for its inhibition. CRISPR-mediated *Sik3* deletion in brown adipocytes increases type IIa histone deacetylase (HDAC) activity and enhances the expression of genes involved in thermogenesis such as *Ucp1*, *Pgc1α*, and mitochondrial OXPHOS complex protein. We further show that HDAC4 interacts with PGC1α after βAR stimulation and reduces lysine acetylation in PGC1α. Finally, a SIK inhibitor well-tolerated *in vivo* (YKL-05-099) can stimulate the expression of thermogenesis-related genes and browning of mouse subcutaneous adipose tissue.

**Conclusions:** Taken together, our data reveal that SIK3, with the possible contribution of other SIKs, functions as a phosphorylation switch for β-adrenergic activation to drive the adipose tissue thermogenic program and indicates that more work to understand the role of the SIKs is warranted. Our findings also suggest that maneuvers targeting SIKs could be beneficial for obesity and related cardiometabolic disease.

## Highlights

- SIK inhibition promotes adipocyte thermogenic gene expression and adipose tissue browning.
- SIK3 interacts with RAPTOR and is phosphorylated by mTOR.
- The regulatory PKA domain is crucial for SIK3 inhibition to drive *Ucp1* expression.
- SIK3 deletion increases type IIa HDAC activity and the thermogenic gene program.
- HDAC4 interacts with PGC1α and decreases the lysine acetylation in PGC1α.

## 1. INTRODUCTION

Adipose tissue plays an essential role in metabolic homeostasis. The white adipose tissue (WAT) functions as a reservoir to store excess calories in the form of lipids, while the brown adipose tissue (BAT) can actively consume glucose and fatty acids and dissipate them as heat by uncoupled respiration through the highly specialized mitochondrial uncoupling protein 1 (UCP1). Brown adipocytes are huge consumers of glucose in order to make fatty acids that are further directed towards oxidation to fuel cold-induced thermogenesis [1]. UCP1-positive adipocytes can be induced in WAT such as subcutaneous inguinal WAT (iWAT) in rodents after cold exposure or treatment with selective agonists for the β_3_-adrenergic receptor (β_3_AR) – a process termed adipose tissue browning [2–7]. Functional BAT has also been identified in adult humans [8–10] and its activity is reported to be associated with an improved cardiometabolic health profile [11]. As a potent stimulator of adipose browning, cold temperature promotes the release of norepinephrine in adipose tissue, which acts through the βAR-cAMP-PKA signaling cascade to drive the expression of a panel of thermogenic and mitochondrial biogenesis genes, including UCP1 and peroxisome proliferator-activated receptor gamma (PPARγ) coactivator-1α (PGC1α) (reviewed in [12; 13]). As an important regulator of the adipocyte thermogenic program, PGC1α activity is regulated by several mechanisms of post-translational modifications such as phosphorylation [14; 15] and reversible acetylation [16–19].

Salt-inducible kinases (SIKs) are members of the AMP-activated protein kinase (AMPK) family. The three SIK isoforms, SIK1, SIK2, and SIK3 are widely expressed and shown to be important regulators of hormonal signaling and metabolism in various tissues [20–23]. Activation of SIKs depends on the phosphorylation in the activation domain by the upstream kinase liver kinase B1 (LKB1), while phosphorylation of SIKs by PKA facilitates their interaction with 14-3-3 proteins and results in SIK inhibition [24]. Several SIK substrates have been identified, such as type IIa histone deacetylases (HDACs) and cAMP response element-binding protein regulated transcription coactivators (CRTCs) [20; 21]. Phosphorylation of these substrates by SIKs leads to their cytoplasmic retention and thus functional inactivation. When SIKs are inhibited by PKA phosphorylation, the SIK substrates will be dephosphorylated, and they can be translocated to the nucleus to regulate target gene expression.

Previous work from our lab demonstrated that activation of the mechanistic target of rapamycin complex 1 (mTORC1) by PKA is essential for β-adrenergic stimulation of adipose tissue browning [25]. Pharmacological mTORC1 inhibition by rapamycin blunts adipose tissue browning in response to the β_3_AR-selective agonist CL-316,243 (CL) treatment, and adipocyte-specific deletion of *Raptor*, the defining component of mTORC1, impairs brown/beige adipocyte expansion during cold challenge [25]. Insulin signaling can also activate mTORC1 through a cascade involving Protein Kinase B (AKT), to promote lipogenesis and suppress lipolysis, a metabolic effect that is antagonized by βAR stimulation [26; 27]. In this study, we hypothesized that PKA activation of mTORC1 regulates the thermogenic gene program in brown adipocytes through the action of specific downstream substrates. We employed a phosphoproteomics approach of Stable Isotope Labeling by/with Amino acids in Cell culture (SILAC) [28] to profile the phosphoproteins in brown adipocytes in response to βAR agonist stimulation. We identified SIK3 as a novel rapamycin-sensitive mTORC1 substrate and further investigated the effects of SIK3 deficiency or SIK inhibition on thermogenic gene expression in brown adipocytes and mouse adipose tissues.

## 2. MATERIALS AND METHODS

### 2.1. Reagents and antibodies

SIK inhibitors HG-9-91-01 (HG, 101147), YKL-05-099 (YKL, 15776), pterosin B (PTB, N1570), type IIa HDAC inhibitors TMP195 (18361), TMP269 (18360) were obtained from MedChemExpress. Isoproterenol (Iso) (I6504), H-89 (B1427), puromycin (P8833), nicotinamide (NAM) (N0636), T3 (T2877), indomethacin (I7378), dexamethasone (D4902), 3-isobutyl-1-methylxanthine (IBMX, I5879), D-(+)-glucose (G7528), dimethyl sulfoxide (DMSO, D8418), bovine serum albumin (BSA) (A7906) were obtained from Sigma-Aldrich. Non-fat dry milk was obtained from RPI Corporation (M17200500). *cOmplete* mini protease inhibitor (04693124001) and *PhosSTOP* phosphatase inhibitor (04906845001) cocktails were obtained from Roche Diagnostics. Rapamycin (Rapa) was from LC Laboratories (R-5000). The β_3_AR-selective agonist CL-316,243 (CL) was a gift from Dr. Elliott Danforth Jr. (American Cyanamid Co., Pearl River, New York, USA).

Dulbecco’s modified Eagle’s medium (DMEM) (11995065), DMEM/F12 (11330032), Penicillin-Streptomycin (15140122), insulin (Ins) (12585014), L-glutamine (25030081), and pyruvate (11360070) were obtained from Gibco. Rosiglitazone (Avandia) was obtained from GlaxoSmithKline. Fetal bovine serum (FBS) was obtained from CPS Serum (FBS-500-HI) or Atlas Biologicals (EF-0500-A). Plasmocin was obtained from InvivoGene (ant-mpp). Polyethylenimine (PEI) was obtained from Polysciences (#23966), Type I collagenase was obtained from Worthington Biochemical (LS004196). Fatty acid free BSA was obtained from CalBioChem (#126575). Polyethylene glycol 300 (PEG300) was obtained from Spectrum Chemical (P0108). Tween-80 was obtained from TCI Chemicals (T2533).

Antibodies directed against SIK3 (#39477, 1:1000), Myc (#2278 and #2276, 1:1000) and acetylated lysine (#9441, 1:1000) were obtained from Cell Signaling Technology (CST). Antibodies against UCP1 (ab23841, 1:1000) and mitochondrial OXPHOS cocktail (ab110413, 1:1000) was obtained from Abcam. Antibody against PGC1α (A11971 and A19674, 1:1000) was obtained from ABclonal. Antibody against COXIV (11242-1-AP, 1:2000), β-actin (20536-I-AP, 1:2000) and GAPDH (10494-I-AP, 1:2000) were obtained from ProteinTech. Antibody against FLAG (A9469 and F1804, 1:2000), rabbit immunoglobulin G (IgG) (A3687, 1:20,000), mouse IgG (A3562, 1:20,000), and anti-FLAG M2 affinity beads (A2220) were obtained from Sigma-Aldrich.

### 2.2. Plasmids and constructs

The RAPTOR-Myc (1859), HDAC4-Flag (13821), PGC1α-Flag (1026), PGC1α-Myc (24925), HDAC4 (21096), GCN5 (63706), pLenti-Puro-sgSIK3-C (138683), pLenti-Puro-sgNT (138681), pLenti-Puro-GFP (17448), pLKO.1-Puro (8453), pLKO.1-Puro-scramble shRNA (1864) plasmids were obtained from Addgene. The NAA10-Myc plasmid was constructed by subcloning of a Myc-His tag to replace the Myc-DDK tag in the hNAA10-Myc-DDK plasmid (RC201354, OriGene). The SIK3-Flag plasmid was constructed by subcloning of a 3xFlag tag to replace the Myc-DDK tag in the mSIK3-Myc-DDK plasmid (MR211912, OriGene). SIK3-Flag-ΔPKA (ΔPKA) was generated by BclI-digestion and relegation of SIK3-Flag plasmid. SIK3-Flag-ΔKD/PKA (ΔKD/PKA) was generated by NheI-digestion and relegation of SIK3-Flag plasmid. SIK3-Flag-S884A and S884D plasmids were generated by introducing site-directed mutations into SIK3-Flag plasmid using the QuikChange II XL site-directed mutagenesis kit according to manufacturer’s instruction (Agilent). Lenti-Puro-SIK3-Flag plasmid was constructed by subcloning the “mSIK3-3xFlag” coding sequences (CDS) from SIK3-Flag plasmids to replace the GFP CDS in the pLenti-Puro-GFP plasmid. The Lenti-Puro-SIK3-Flag plasmids with ΔPKA, ΔKD/PKA, S884A, or S884D mutations/deletions were constructed with the same subcloning procedure. The pLenti-Puro derived plasmids are used for over-expression studies in **Figure 2H** and **3G**. The shRNA plasmids for mouse *Pgc1α*, *Sik1*, *Sik2*, *Sik3*, *Hdac4*, *Hdac5*, and *Hdac7* were constructed by subcloning the annealed short hairpin oligos to the pLKO.1-Puro vector as previously described [29; 30]. The sequences of the short hairpins are: *mPgc1α*: AGACTATTGAGCGAACCTTAA; *mSik1:* TGTAGTGGTTGGCGTTGTTTC; *mSik2*: GTCTCAGCTGCAAGCATATTT; *mSik3*: CGCACGGAAGTTATGGAAGAT; *mHdac4*: GGTACAATCTCTCTGCCAAAT; *mHdac5*: GGCTCAGACAGGTGAGAAAGA; *mHdac7*: GGGTCGATACTGACACCATCT. Plasmids are available on request as listed in **Supplementary Table 1**.

### 2.3. Cell lines

HEK293FT cells were from Invitrogen. HIB-1B brown adipocyte cells were a gift from Dr. Bruce Spiegelman [31]. HEK293FT and HIB-1B cells were cultured in Growth Medium (DMEM with 10% FBS, 100 U/ml Penicillin, 100 μg/ml Streptomycin) supplemented with 25 μg/ml plasmocin. HIB-1B cells were induced for differentiation in Growth Medium with 1 μM rosiglitazone for three to four days, and then maintained in Growth Medium until experiments. Primary brown adipocyte cultures were prepared as previously described [32; 33]. Briefly, the stromal vascular cells of the interscapular BAT were isolated via collagenase digestion of BAT from 6-8 weeks-old wild-type mice, cultured in DMEM/F12 medium with 15% FBS, and split into 6-well plates. When reach 95% confluency, cells were induced for differentiation for three days in DMEM/F12 medium with 10% FBS supplemented with a cocktail of 5 μg/ml insulin, 1 μM T3, 1 μM rosiglitazone, 125 μM indomethacin, 1 μM dexamethasone, and 0.5 mM IBMX. Cells were then maintained in DMEM/F12 medium with 10% FBS, 5μg/ml insulin, 1 μM T3, and 1 μM rosiglitazone for additional 3-4 days until they were ready for experiments.

### 2.4. shRNA knockdown and SIK3 over-expression

HIB-1B cells were seeded in 6-well plates at a density of 5 x 10^5^ cells/well and reached 90% confluence when ready for transfection the following day. Cells were transfected with 2.5 μg/well shRNA knockdown (pLKO.1-Puro) or SIK3 expression (pLenti-Puro-3xFlag) plasmids (Supplementary Table 1) by PEI reagent as previously described [34]. Six-hours after transfection, cells were induced for differentiation in Growth Medium with 1 μM rosiglitazone. Twenty-four hours after transfection, 3 μg/ml puromycin were added to the differentiation medium for drug selection for 2 days. Three or four days after differentiation, cells were replenished with fresh Growth Medium and allowed to recover at least for 3 hours before treatments for gene expression analysis.

### 2.5. CRISPR gene editing

HIB-1B cells were plated in 6-well plates at a density of 5 x 10^5^ cells/well and reached 90% confluence when they were ready for transfection the following day. Cells were transfected with 2.5 μg/well pLenti-Puro-sgSIK3-C (SC) or pLenti-Puro-sgNT plasmids by PEI reagent as previously described [34]. Six-hours after transfection, cells were replenished with Growth Medium. Twenty-four hours after transfection, 2 μg/ml puromycin were added to the medium for drug selection for 3 days. Cells were then supplemented with fresh Growth Medium, allowed to recover for 9 hours, trypsinized and replated in 96-well plate at a density of 1.6 cell/100ul/well. Single-cell clones were passaged and cryo-stored in liquid nitrogen. The genomic DNA were extracted from each clone with PureLink Genomic DNA Mini Kit (Invitrogen) and PCR amplified with the forward primer TCCCCTTCAGCTCTGTTCAG (mSik3-sgC-F1) and reverse primer AAGTGAAGTCTGGAGTGGGG (mSik3-sgC-R1). The PCR products were subcloned to T-vector (Promega) and sequenced. Two clones with frame-shifted mutations in *Sik3* gene were validated for the efficient knockout of SIK3 protein and considered as *Sik3*^-/-^ brown adipocytes (SC19 and SC13). A pool of twelve single clones derived from the mock transfection of pLenti-Puro-sgNT plasmid were used as non-targeting control (NTC).

### 2.6. SILAC

The immortalized murine brown adipose cells were generated in the lab of Dr. C. Ronald Kahn, originally derived from BAT of newborn mice, and cultured as previously described [35]. For SILAC study, shown in **Figure 1A**, the cells were cultured and labeled with light (Arg0, Lys0), middle (Arg6, Lys4) and heavy (Arg10, Lys8) amino acids as previously described [28], and then cultured for 12-16 h with media containing 1% Fetal Calf Serum (FCS). The labeled cells were then treated with Ins (10 nM), Iso (1 µM) and/or Rapa (100 nM). Rapa was pre-incubated for 30 min followed by the treatment with Ins or Iso for 15 min. The middle-labeled cells were used as an internal standard. After treatment, cells were lysed with 500 µl 6 M guanidinium chloride. The light, middle and heavy lysates were mixed 1:1:1. Proteins were precipitated over night with 100% acetone followed by in-solution LysC digestion and trypsin digestion overnight. After desalting, the sample peptides were separated with high pH-HPLC fractionation. The 12 pooled fractions were extracted two times with TiO2, and samples were put onto C8 stage tips (High-Select™ TiO2 Phosphopeptide Enrichment Kit; ThermoFisher), eluted, and then analyzed with a 90 min LC-MS/MS gradient at the Q Exactive mass spectrometer. The data was evaluated with MaxQuant and Perseus software. Every experiment was done in triplicate.

**Figure 1:**
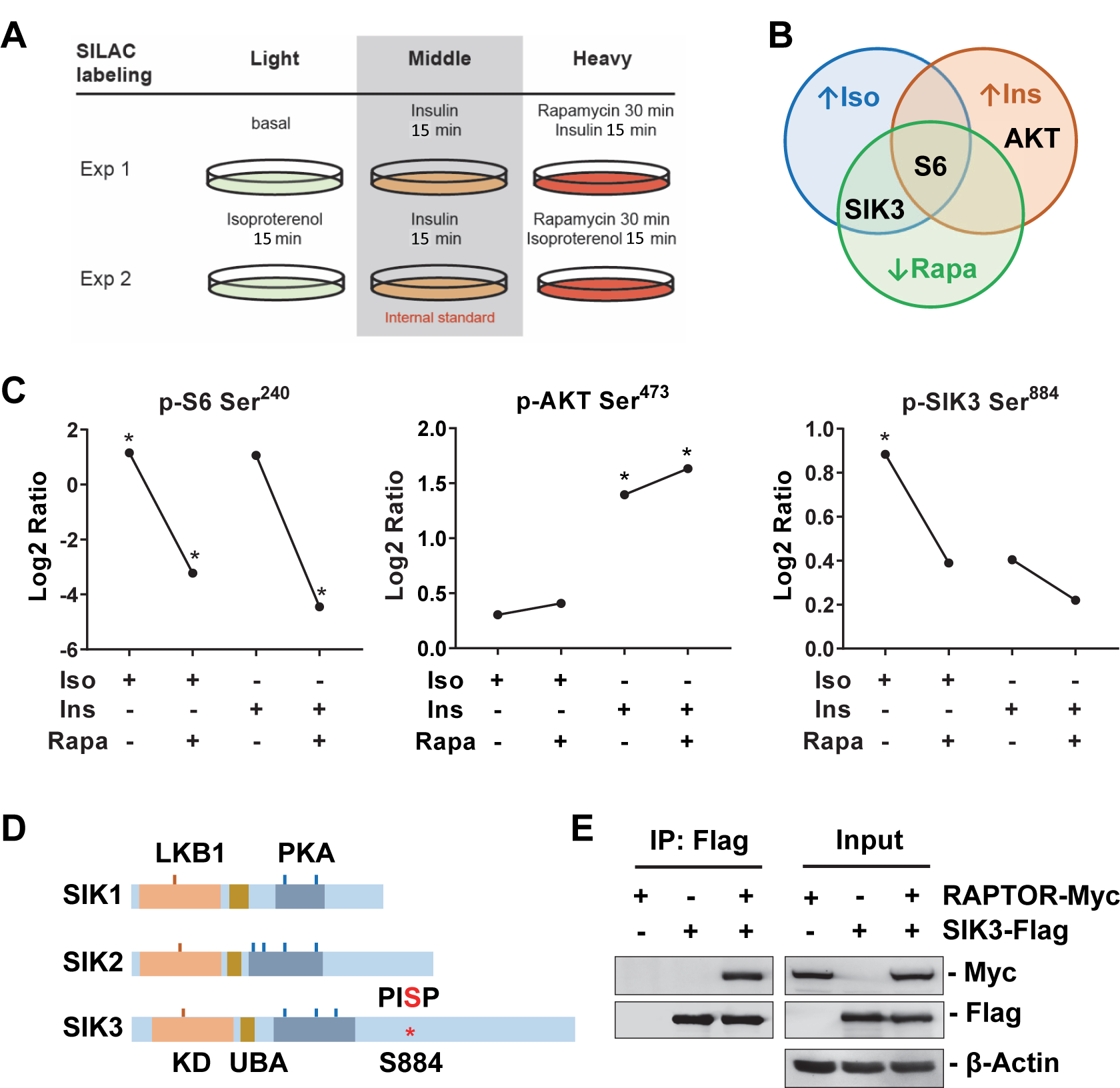
SILAC studies identified SIK3 as a new mTORC1 substrate. (A) Murine brown adipose cell labeling and treatment for SILAC analysis. Isotope labeled cells that received rapamycin were pretreated for 30 min, and then treated with isoproterenol or insulin for 15 min. Cell lysates were processed and subjected to liquid chromatography and mass spectrometry analysis to profile the phosphorylated peptides. (B) Venn diagram illustration of the representative protein whose phosphorylation is increased by isoproterenol (Iso, blue) or insulin (Ins, orange) and blocked by rapamycin (Rapa, green). (C) The phosphorylation of S6 Ser^240^, AKT Ser^473^ and SIK3 Ser^884^ in response to Iso, Ins and Rapa treatment. Data are presented as log2 fold change compared to the vehicle treated samples. *p<0.05. (D) SIK protein structure and phosphorylation sites. The LKB1 (Liver kinase B1, orange), PKA (blue), and mTORC1 (S884, red) phosphorylation sites are indicated. KD: kinase domain. UBA: ubiquitin-associated domain. (E) HEK293FT cells were transfected with SIK3-Flag and/or RAPTOR-Myc plasmids as indicated. Protein lysates were immunoprecipitated with anti-Flag beads and blotted with indicated antibodies.

### 2.7. Cell mitochondrial stress test

Control NTC (2,000 cells/well), *Sik3*^-/-^ SC19 (2,500 cells/well), and SC13 (5,000 cells/well) cells were seeded in Agilent Seahorse XF96 microplates, differentiated in Growth Medium with 1 μM rosiglitazone for three days. For Seahorse XF Cell Mito Stress test (Agilent, 103015-100), cells were incubated in Seahorse XF assay DMEM medium (Agilent, 103575-100, pH 7.4) supplemented with 10 mM glucose, 2 mM glutamine, and 1 mM pyruvate. Cellular respiration rate was determined at baseline and after the sequential addition of ATP synthase inhibitor oligomycin (Oligo, 2 μM), mitochondrial uncoupler Carbonyl cyanide p-trifluoro-methoxyphenyl hydrazone (FCCP, 0.5 μM), and respiration chain complex I/III inhibitors rotenone/antimycin A (Rot/AA, 0.5 μM) on a Seahorse extracellular flux XF96 analyzer according to manufacturer’s instructions. After Seahorse assay, cell nuclei were stained with 9 μM Hoechst 33342 (Invitrogen, H3570) for 10 min, imaged with ImageXpress (Molecular Device), and counted using the multi wavelength cell scoring protocol of MetaXpress software (Molecular Device). The respiration rate was normalized as oxygen consumption rate per 10,000 cells as shown in **Figure 3D**. The basal, proton leak, maximal, and non-mitochondrial respiration were calculated according to manufacturer’s instructions.

**Figure 2:**
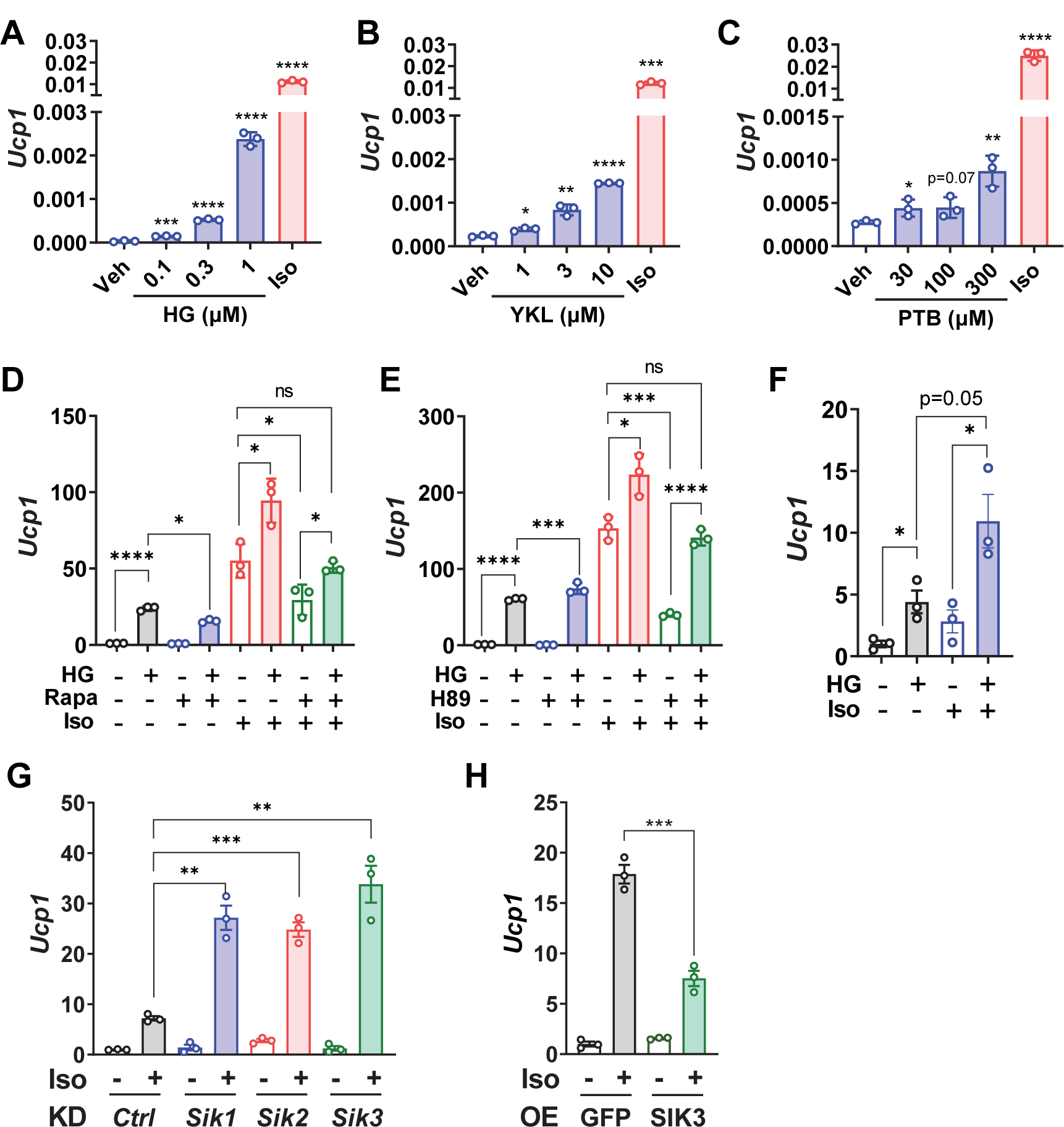
SIK inhibition stimulates thermogenic gene expression in brown adipocytes. (A-C) SIK inhibitors increase *Ucp1* mRNA expression dose-dependently. HIB-1B cells were treated with indicated amount of SIK inhibitors HG-9-91-01 (HG), YKL-05-099 (YKL), and pterosin B (PTB) for 6 hrs, or with 100nM Iso for 3 hrs. *Ucp1* mRNA level was determined by QPCR. Data show relative expression levels normalized to the internal control gene 36B4. Student’s t-test, compared with vehicle, *p<0.05, **p<0.01, ***p<0.001, ****<0.0001. (D-E) SIK inhibitor restores *Ucp1* expression after blockades of mTORC1 (by Rapa) or PKA (by H89). HIB-1B cells were pretreated with 1μM HG (3 hrs), 100nM Rapa (16 hrs) or 20μM H89 (3 hrs) and then treated with 100nM Iso for 3 hrs. Student’s t-test, *p<0.05, ****<0.0001, ns: no statistical difference. (F) Primary mouse brown adipocytes were pretreated with 1μM HG for 3 hrs and then treated with 100nM Iso for 6 hrs. Student’s t-test, *p<0.05. (G) shRNA knockdown (KD) of either *Sik1*, *Sik2*, or *Sik3* increases *Ucp1* expression in HIB-1B brown adipocytes. (H) Overexpression (OE) of SIK3 reduces *Ucp1* expression in HIB-1B brown adipocytes. After transfection of shRNA or SIK3 plasmids, HIB-1B cells were differentiated for three days, replenished with fresh medium for 3 hrs, and treated with 100nM Iso for 3 hrs. Student’s t-test, *p<0.05, **p<0.01, ***p<0.001, ****<0.0001, ns: no statistical difference.

**Figure 3:**
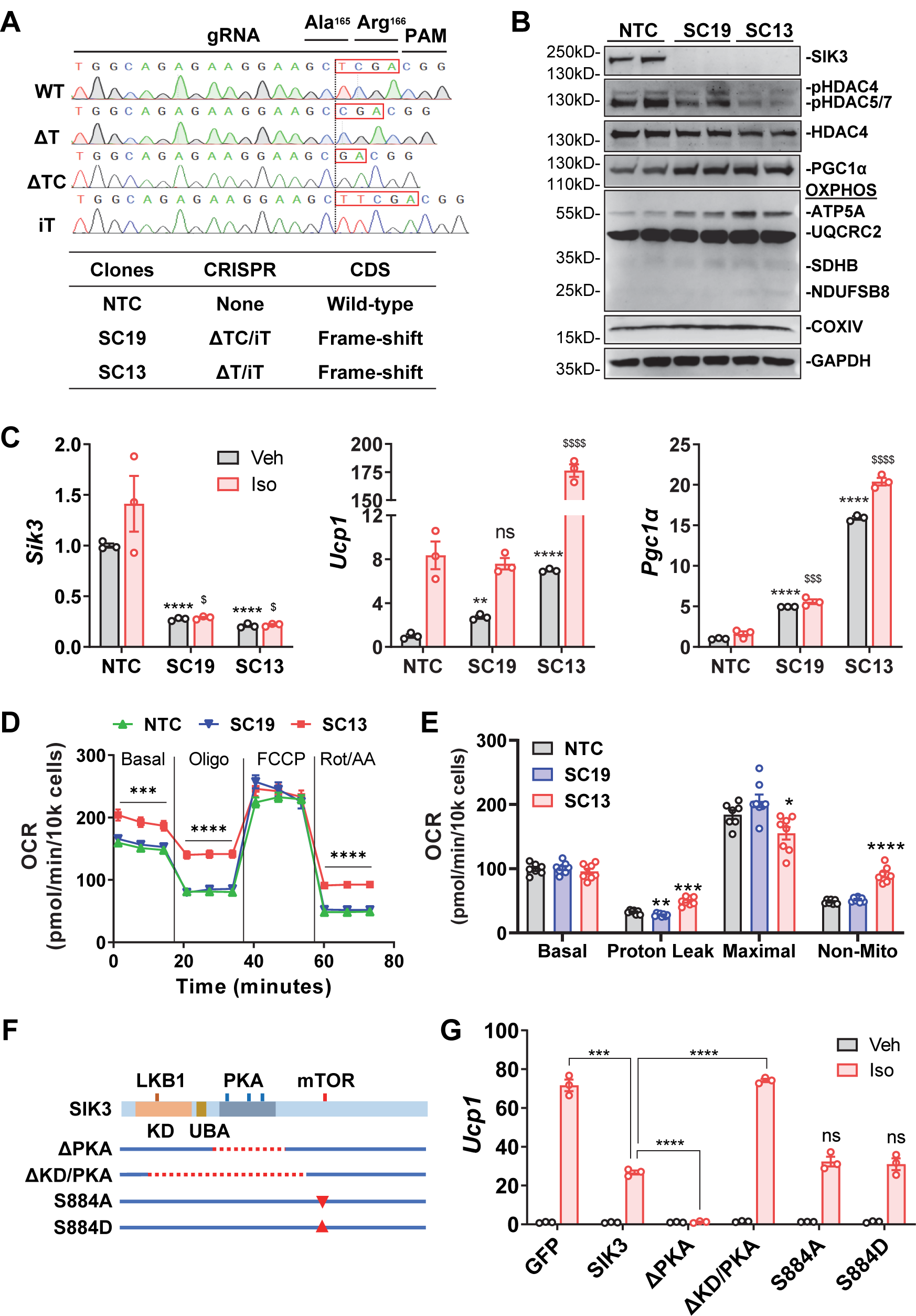
*Sik3* deletion in brown adipocytes increases thermogenic gene expression. (A) Sequence of wild-type (WT) and CRISPR-edited *Sik3* mutations, the guide RNA (gRNA), and the protospacer adjacent motif (PAM) are shown (upper). The wild-type non-targeting control (NTC), and two CRISPR-edited HIB-1B clones with frameshifted *Sik3* mutations (SC19 and SC13) were generated (lower). (B) Western blot analysis of SIK3, type IIa HDACs, PGC1α and mitochondrial OXPHOS proteins ATP5A, UQCRC2, SDHB, NDUFSB8, COXIV in control (NTC) and *Sik3*^-/-^ (SC19 and SC13) brown adipocytes. p-HDAC4 Ser246, p-HDAC5 Ser259, p-HDAC7 Ser155. OXPHOS: oxidative phosphorylation complex. (C) Expression of *Sik3*, *Ucp1* and *Pgc1α* mRNA in control (NTC) and *Sik3*^-/-^ (SC19 and SC13) brown adipocytes. Student’s t-test, **p<0.01, ***p<0.001, ****p<0.0001 vs NTC vehicle (Veh); ^$^p<0.01, ^$$$^p<0.001, ^$$$$^p<0.0001 vs NTC Iso. (D-E) Cell mitochondrial stress test of control (NTC) and *Sik3*-/-(SC19 and SC13) brown adipocytes. Oligo: oligomycin; FCCP: carbonyl cyanide p-trifluoro-methoxyphenyl hydrazone; Rot/AA: rotenone and antimycin A. The basal, proton leak, maximal, and non-mitochondrial respiration were presented. Student’s t-test, *p<0.05, **p<0.01, ***p<0.001, ****p<0.0001 vs NTC, (F) SIK3 deletion/mutation constructs. ΔPKA: Δ432-805, PKA phosphorylation domain. ΔKD/PKA: Δ147-850, kinase domain (KD), ubiquitin-associated domain (UBA), and PKA phosphorylation domain. S884A: Ser884 to Ala phosphorylation-resistant mutation. S884D: Ser884 to Asp phosphorylation-mimetic mutation. (G) Expression of *Ucp1* mRNA in *Sik3*^-/-^ brown adipocytes (SC19) transfected with GFP, SIK3, or SIK3 deletion/mutation plasmids. At day 3 of differentiation, cells were recovered in fresh medium for 3 hrs and treated with vehicle (Veh) or 100nM Iso for 3 hrs. Student’s t-test, ***p<0.001, ****p<0.0001; ns: no statistical difference vs SIK3 Iso.

### 2.8. Mouse YKL treatment

Twelve to thirteen-weeks-old male C57BL/6J (Jackson Laboratory) mice were administrated vehicle (5% DMSO, 40% PEG300, 5% Tween-80, 50% Saline) or YKL at doses of 10 or 20 mg/kg/day via intraperitoneal (i.p.) injection for 10 days. As a positive control, a group of mice were administrated β_3_AR selective agonist CL at a dose of 1 mg/kg/day i.p. for 5 days. The food intake of each group was measured manually every day. All animal studies were approved by the Institutional Animal Care and Use Committee of Vanderbilt University Medical Center and in accordance with the NIH Guide for the Care and Use of Laboratory Animals.

### 2.9. Histology and morphological segmentation analysis of adipocyte size

Adipose tissues were fixed in 10% neutral buffered formaldehyde overnight, routinely processed for paraffin section, and stained with hematoxylin and eosin (H&E) or immunohistochemically stained for UCP1 (ab10983, Abcam). Histology was performed by the Vanderbilt Translational Pathology Shared Resource Core. Whole slides were imaged at 20X magnification to a resolution of 0.5 µm/pixel using a high throughput Leica SCN400 Slide Scanner automated digital image system by the Digital Histology Shared Resource Core at Vanderbilt University Medical Center (www.mc.vanderbilt.edu/dhsr). For adipocyte size analysis, ten H&E stained images of 1 mm^2^ (4,000,000 pixel^2^) were randomly extracted from the entire adipose tissue image using Aperio ImageScope (Leica Biosystems Imaging, Deer Park, IL) and analyzed with the MorphoLibJ collection of mathematical morphology methods and plugins using the image processing package FIJI [36]. The image color was converted into binary with the color threshold in the range of 0 to 35 to ensure that cells were clearly visible and distinguishable. Adipocyte size (pixel^2^) and total adipocyte counts were determined with a particle size parameter in the range of 50 to 6,000 pixel^2^ and converted into x10 μM^2^ as shown in **Figure 5I-5K**.

### 2.10. Quantitative PCR (QPCR)

RNA was extracted with TRIzol (Invitrogen), purified with RNeasy Mini columns (Qiagen) or Quick-RNA columns (Zymo Research), and reversed transcribed with High-Capacity cDNA Reverse Transcription Kits (Applied Biosystems). QPCR was performed with *PowerUp* SYBR Green Master Mix (Life Technologies) per manufacturer’s instruction. QPCR data were analyzed using the ΔΔCt method, normalized to the internal control gene 36B4, and presented as fold change relative to the control group unless otherwise specified. Primer sequences are listed in **Supplementary Table 2**.

### 2.11. Western blot

Protein was extracted as previously described [33]. In brief, adipose tissues or cells were lysed and sonicated in Protein Lysis Buffer (25 mM HEPES, pH 7.4, 150 mM NaCl, 5 mM EDTA, 5 mM EGTA, 5 mM glycerophosphate, 0.9% Triton X-100, 0.1% IGEPAL, 5 mM sodium pyrophosphate, 10% glycerol, 1x protease inhibitor and 1x phosphatase inhibitor cocktails). For Western blot, 30 to 40 μg of protein was resolved by 10% SDS-polyacrylamide gel electrophoresis (PAGE), transferred to nitrocellulose membranes (Bio-Rad), incubated overnight at 4 °C with primary antibodies in blocking buffer (5% non-fat dry milk or BSA in Tris-buffered saline, 0.1% Tween-20), and followed by incubation with alkaline phosphatase–conjugated secondary antibody for 1 hour at room temperature. Blots were incubated with Amersham ECF substrate (Cytiva, RPN5785) and images were acquired by Bio-Rad digital ChemiDoc MP with IR (VUMC Molecular Cell Biology Resource).

### 2.12. Immunoprecipitation

HEK293FT cells were seeded in 6-well plates at a density of 6 x 10^5^ cells/well, transfected with 2.5 μg/well of indicated plasmids on the following day, and harvested for protein lysate in Protein Lysis Buffer 24 hours after transfection. Cells were treated with 1 μM Iso for 6 hours as indicated before harvesting protein lysates. For immunoprecipitation, 300-500 μg protein lysates were incubated with 15 μl of anti-FLAG M2 Affinity beads in Protein Lysis Buffer at 4 °C for 2 hour s. Affinity beads were washed three times and eluted with 20 μl 2x NuPAGE LDS sample buffer (Invitrogen, NP0008) by incubation at 95 °C for 5 min. Protein eluent were subjected to SDS - PAGE and Western blot analysis as described above.

### 2.13. PGC1α acetylation

HEK293FT cells were plated in 10-cm plate at a density of 3 x 10^6^ cells/plate, and on the following day transfected with 15 μg plasmids/plate, including PGC1α-Flag, GCN5, NAA10-Myc, with vector or HDAC4. Cells were replenished with fresh medium 6 hours after transfection, treated with 5 mM nicotinamide overnight and then treated with 1 μM Iso for 6 hours on the following day, and lysed in 1x Cell Lysis Buffer (CST, #9803) supplemented with 1x protease inhibitor and 1x phosphatase inhibitor cocktail. For Western blot analysis of PGC1α acetylation, 500 μg protein lysate were used for immunoprecipitation and blotted with anti-acetylated lysine antibody as described above.

### 2.14. Statistical analysis

GraphPad Prism and Microsoft Excel were used for statistical analysis. All data were presented as means ± SEM. Unpaired two-tailed Student’s t-tests were used to determine the differences between groups. Statistical significance was defined as p < 0.05.

## 3. RESULTS

### 3.1. SILAC studies identified SIK3 as a novel mTORC1 substrate in murine brown adipocytes

We employed a phosphoproteomic approach of Stable Isotope Labeling by/with Amino acids in Cell culture (SILAC) [28] to profile the global protein phosphorylation in murine brown adipocytes after either βAR agonist isoproterenol (Iso) or insulin (Ins) stimulation, in combination with or without the mTORC1 inhibitor rapamycin (Rapa) (**Figure 1A**). As a well-established substrate of the mTOR signaling pathway, phosphorylation of Ser^240^ in ribosomal protein S6 was increased both by Iso and Ins and blocked by rapamycin, while phosphorylation of Ser^473^ in AKT was increased only by Ins and unaffected by Iso or Rapa, as its activation is upstream of mTORC1 (**Figure 1B** and **1C**). These results mirror our previous findings [25]. Interestingly, among several other proteins, the phosphorylation of SIK3 on Ser^884^ (S^884^) was increased by Iso, blocked by Rapa, but unaffected by Ins (**Figure 1B** and **1C**), suggesting a mTORC1-dependent effect. The S^884^ site (PISP) conforms to identified mTOR phosphorylation sites [37] and is found in the C-terminal regulatory domains of SIK3 but does not exist in the other two salt-inducible kinases SIK1 or SIK2 (**Figure 1D**). Subsequent experiments showed that SIK3 can be immunoprecipitated with RAPTOR, the defining component of mTORC1 and which is known to present substrates to the mTOR kinase [38; 39], indicating the physical interaction of SIK3 with mTORC1 (**Figure 1E**).

### 3.2. SIK inhibition stimulates *Ucp1* gene expression in HIB-1B brown adipocytes

To understand the role of SIK3 in the brown adipocyte thermogenic gene program, we next examined the effect of pharmacological SIK inhibition in HIB-1B murine brown adipocytes. Several compounds have been reported to inhibit SIK activities, including the pan-SIK inhibitors HG-9-91-01 (HG) [40; 41], YKL-05-099 (YKL) [42] and the SIK3-selective inhibitor pterosin B (PTB) [43; 44]. Our results showed that all three compounds increase basal expression of the *Ucp1* gene in HIB-1B cells in a dose-dependent manner (**Figure 2A-2C**). While the βAR agonist Iso is a more potent stimulator, the point here is that even in the absence of Iso there is nevertheless an increase in basal *Ucp1* expression in conditions in which βAR activity is low.

Because inhibition of PKA or mTORC1 prevents βAR agonist stimulated *Ucp1* expression and adipose browning [25], we next asked whether SIK inhibition could override the effects of PKA or mTORC1 inhibitors. Pretreatment with HG increased *Ucp1* expression under basal unstimulated conditions and potentiated the response to Iso (**Figure 2D** and **2E**). HG also restored *Ucp1* expression after either mTORC1 inhibition by Rapa (**Figure 2D**) or PKA inhibition by H89 (**Figure 2E**). This would place SIK inhibition as a step downstream of the PKA-mTORC1 signaling cascades after β-adrenergic stimulation. In line with the results in HIB-1B brown adipocytes, HG treatment also increased *Ucp1* expression under basal conditions and further augmented its expression in response to Iso stimulation in primary mouse brown adipocytes (**Figure 2F**).

Since this effect of SIK inhibition on *Ucp1* gene expression is not SIK isoform-specific, we next sought to dissect the role of SIK3 by specific manipulation of SIK3 gene expression in HIB-1B cells. Short hairpin RNA (shRNA) knockdown of either *Sik1*, *Sik2*, or *Sik3* increased *Ucp1* gene expression in HIB-1B cells after Iso stimulation (**Figure 2G**), indicating that SIKs might play redundant roles in regulating *Ucp1* expression in brown adipocytes. In addition, over-expression of SIK3 in HIB-1B cells suppressed *Ucp1* expression (**Figure 2H**), supporting the notion that SIK3 functions as a suppressor of brown adipocyte thermogenic gene program.

### 3.3. SIK3 deletion in brown adipocytes increases thermogenic gene expression

We next utilized CRIPSR/Cas9 gene editing to specifically knockout the *Sik3* gene in HIB-1B brown adipocytes using a previously established guide RNA (gRNA) expression construct [45]. Sequencing results showed that three major mutations were generated in the *Sik3* gene: a single-nucleotide deletion (ΔT), a two-nucleotide deletion (ΔTC) and a single-nucleotide insertion (iT) (**Figure 3A**). Each of these mutations will generate transcripts with frameshifts from the Ala^165^ (A^165^) residue, resulting in truncated proteins due to premature stop codons (**Figure 3A**). Two clones, SC19 and SC13, were identified to bear two frame-shift mutations and considered *Sik3*^-/-^ brown adipocytes (**Figure 3A**). Western blot analysis showed that SIK3 protein was completely absent in these *Sik3*^-/-^ adipocytes but not in the non-targeting control (NTC) cells (**Figure 3B**). Interestingly, the phosphorylation of SIK3 substrates HDAC4, HDAC5 and HDAC7 were decreased, suggesting enhanced type IIa HDAC activities in *Sik3*^-/-^ adipocytes (**Figure 3B**). In addition, the expression of PGC1α and certain mitochondrial oxidative phosphorylation (OXPHOS) proteins were increased in *Sik3*^-/-^ adipocytes (**Figure 3B**). Furthermore, QPCR analysis showed that *Sik3* mRNA was significantly reduced in the SC19 and SC13 brown adipocytes (**Figure 3C**), potentially due to nonsense-mediated mRNA decay [46]. Expression of *Ucp1* and *Pgc1α* mRNA were increased in *Sik3*^-/-^ adipocytes both at basal conditions and after Iso stimulation (**Figure 3C**). This data is consistent with the *Sik3* knockdown and over-expression experiments (**Figure 2G** and **2H**). Of note, although the SIK3 protein was absent from both SC19 and SC13 clones, the increases in *Ucp1* and *Pgc1α* mRNA expression were more robust in clone SC13 (**Figure 3C**), possibly due to clonal variations or other potential off-target effects as previously reported [47]. Nevertheless, in both clones the pattern of increased thermogenic gene expression was evident.

We next performed respiration measurements in the *Sik3*^-/-^ adipocytes by a mitochondrial stress test. Results showed that the *Sik3*^-/-^ adipocyte SC13 clone had a higher basal oxygen consumption rate (OCR). In the presence of the ATP synthase inhibitor oligomycin there was more residual respiration (**Figure 3D** and **3E**), indicating a greater degree of uncoupled respiration, also readily observed by the shallower slope, potentially due to the enhanced *Ucp1* expression or function. The *Sik3*^-/-^ clone SC13 also had a higher degree of non-mitochondrial respiration upon inhibition of the respiratory chain complexes I and III by rotenone and antimycin (**Figure 3D** and **3E**). However, the *Sik3*^-/-^ clone SC19 did not exhibit this same elevated OCR and was similar to the NTC control cells. Due to the phenotypic variations between *Sik3*^-/-^ clones SC19 and SC13, characterization of additional CRISPR-edited clones will be needed in future to more comprehensively assess the effect of SIK3 deficiency on cellular respiration in brown adipocytes.

### 3.4. The regulatory PKA phosphorylation domain is essential for SIK3 inhibition to drive thermogenic gene expression

Using our *Sik3*^-/-^ HIB-1B brown adipocytes, we next established a SIK3 re-expression procedure to evaluate the functional domains of SIK3 protein in relation to *Ucp1* gene expression. It is well-established that phosphorylation of SIKs by LKB1 in the N-terminal kinase domain (KD) is essential for SIK activation, while phosphorylation by PKA in the internal regulatory domain results in SIK inhibition, which is no longer able to phosphorylate SIK substrates and excludes them from nuclear accumulation [20; 21]. We constructed expression plasmids with deletions of the PKA phosphorylation domains alone (ΔPKA) or in combination with the kinase domain (ΔKD/PKA) (**Figure 3F**). We also constructed expression plasmids with mutations of the S^884^ site into either phosphorylation-resistant S^884A^, or phosphorylation-mimetic S^884D^ (**Figure 3F**). These SIK3 proteins with deletions or S^884^ mutations were re-expressed in *Sik3*^-/-^ HIB-1B cells (clone SC19), and the effects on *Ucp1* mRNA expression were evaluated. Consistent with our previous experiment (**Figure 2H**), re-expression of wild-type SIK3 protein significantly suppressed *Ucp1* expression after Iso stimulation (**Figure 3G**). More importantly, deletion of the internal PKA regulatory region in SIK3 abrogated PKA phosphorylation and resulted in a constitutively active SIK3 that suppresses *Ucp1* expression regardless of Iso stimulation (**Figure 3G**), suggesting the regulatory PKA phosphorylation domain is essential for SIK3 inhibition that allows thermogenic gene expression. In contrast, deletion of the kinase domain along with the PKA domain in SIK3 impaired its capacity to suppress *Ucp1* expression (**Figure 3G**), suggesting that this inhibitory effect depends on its kinase activity. However, re-expression of either the phosphorylation-resistant SIK3 S^884A^ or the phosphorylation-mimetic SIK3 S^884D^ mutation did not change *Ucp1* expression (**Figure 3G**), indicating that the S^884^ phosphorylation state alone does not alter the function of SIK3 in regulating *Ucp1* gene expression. This finding suggested to us that mTORC1-dependent phosphorylation of SIK3 at S^884^ might affect some other function in the overall βAR-dependent regulation of brown adipocytes that we have not yet identified. An unbiased approach in cells expressing the S^884^ mutants may yield new insight.

### 3.5. HDAC4 is involved in SIK inhibition to promote thermogenic gene expression

Type IIa HDACs are the most well-studied SIK substrates, including HDAC4, HDAC5, and HDAC7 [20; 21]. To determine whether type IIa HDACs are downstream mediators of SIK inhibition to drive *Ucp1* gene expression, we next tested the effect of type IIa HDAC inhibitors on *Ucp1* gene expression in HIB-1B brown adipocytes. Our results show that type IIa HDAC inhibitors TMP195 and TMP269 significantly reduced *Ucp1* gene expression after Iso stimulation (**Figure 4A**). In addition, knockdown of *Hdac4* in HIB-1B brown adipocytes blunted *Ucp1* gene expression upon SIK inhibition by the HG compound (**Figure 4B**). Similarly, knockdown of *Pgc1α*, a key regulator of *Ucp1* expression in brown adipocytes, also abolished *Ucp1* induction after SIK inhibition (**Figure 4C**). Although the phosphorylation of HDAC5 and HDAC7 were also decreased in *Sik3*^-/-^ brown adipocytes (**Figure 3B**), shRNA knockdown of either *Hdac5* or *Hdac7* did not alter *Ucp1* gene expression upon HG treatment (**Figure 4D**). These results suggested that SIK inhibition controls *Ucp1* gene expression potentially through the action of HDAC4.

**Figure 4:**
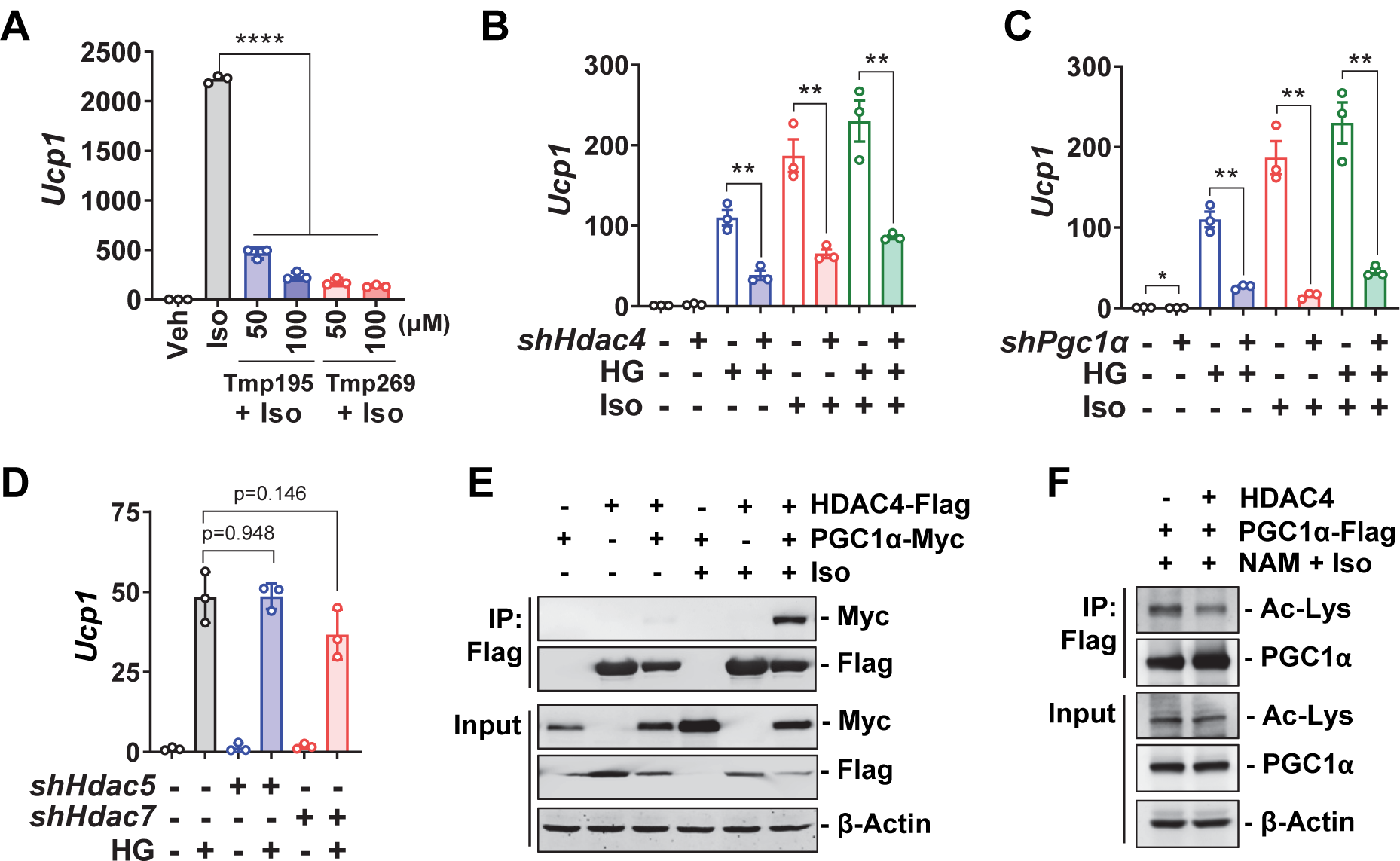
HDAC4 is involved in SIK inhibition to promote thermogenic gene expression. (A) Type IIa HDAC inhibitors blunt Iso-stimulated *Ucp1* expression. At day 3 of differentiation, HIB-1B cells were pretreated with Tmp195 and Tmp269 for 16 hrs and then treated with 100 nM Iso for 3 hrs. Student’s t-test, ****p<0.0001. (B-D) Knockdown of *Hdac4* or *Pgc1α*, but not *Hdac5* or *Hdac7*, blunts the increase of *Ucp1* expression after SIK inhibition. After transfection of shRNA plasmids, HIB-1B cells were differentiated for three days and replenished with fresh medium overnight. On the following day, cells were pretreated with 1μM HG for 3 hrs, and then treated with 100nM Iso for 3 hrs. Student’s t-test, **p<0.01; *p<0.05. (E) HDAC4 interacts with PGC1α. HEK293FT cells were transiently transfected with indicated plasmids and treated with 1 μM Iso for 6 hrs. Protein lysates were immunoprecipitated with anti-Flag antibody-conjugated beads and blotted with indicated antibodies. (F) PGC1α-Flag, GCN5, NAA10-Myc plasmids were co-transfected either with HDAC4 or vector into HEK293FT cells. Cells were treated with 5mM nicotinamide (NAM, SIRT1 inhibitor) overnight followed by 1 μM Iso for 6 hrs. PGC1α were immunoprecipitated with anti-Flag beads and blotted with anti-acetylated lysine (Ac-Lys) or PGC1α antibodies.

It has been shown that deacetylation of PGC1α is required for its optimal coactivator activity [19]. Therefore, we asked whether HDAC4-mediated PGC1α deacetylation could be a potential mechanism by which SIK inhibition promotes *Ucp1* gene expression. In support of this hypothesis, we found that HDAC4 can be co-immunoprecipitated with PGC1α (**Figure 4E**), in line with a previous report [48]. Of note, βAR agonist stimulation is required for the interaction of HDAC4 with PGC1α (**Figure 4E**). This is consistent with previously reports showing that cAMP signaling facilitates the nuclear translocation of HDAC4 [49; 50]. More importantly, co-expression of HDAC4 can reduce the lysine acetylation of PGC1α (**Figure 4F**). We also observed that lysine deacetylation seemed to increase the total PGC1α protein level (**Figure 4F**), indicating that lysine deacetylation might increase PGC1α protein stability. Since the PGC1α antibody recognizes the C-terminal region of the protein, we do not know yet whether the deacetylation in this region will allow it to be better recognized by the antibody or not.

### 3.6. SIK inhibitor YKL stimulates thermogenic gene expression and adipose browning *in vivo*

YKL has been shown to be an effective SIK inhibitor *in vivo* [42] and has been applied to disease models such as osteoporosis and leukemia [29; 51]. Our data (**Figure 2B**) showed that YKL can increase *Ucp1* gene expression in HIB-1B brown adipocyte *in vitro*. We next sought to examine the effect of YKL on adipose tissue browning *in vivo*. Chow diet fed mice housed at room temperature were administered vehicle, YKL, or the β_3_AR agonist CL as a positive control for adipose browning (**Figure 5A**). YKL was delivered in daily doses of 10 and 20 mg/kg bodyweight for 10 days. As shown in **Figure 5B**, while the body weight and tissue morphology appeared normal, the weights of several tissues from mice of the 20 mg group were slightly altered, including the gonadal WAT (gWAT), heart, and spleen (**Figure 5C**). In addition, the daily food intake was significantly reduced in mice of the 20 mg group (**Figure 5D**), but the fasting blood glucose levels were normal in both dosage groups (**Figure 5E**). These data indicated that while the 10 mg dose was well-tolerated, the 20 mg dose might elicit other side effects.

**Figure 5:**
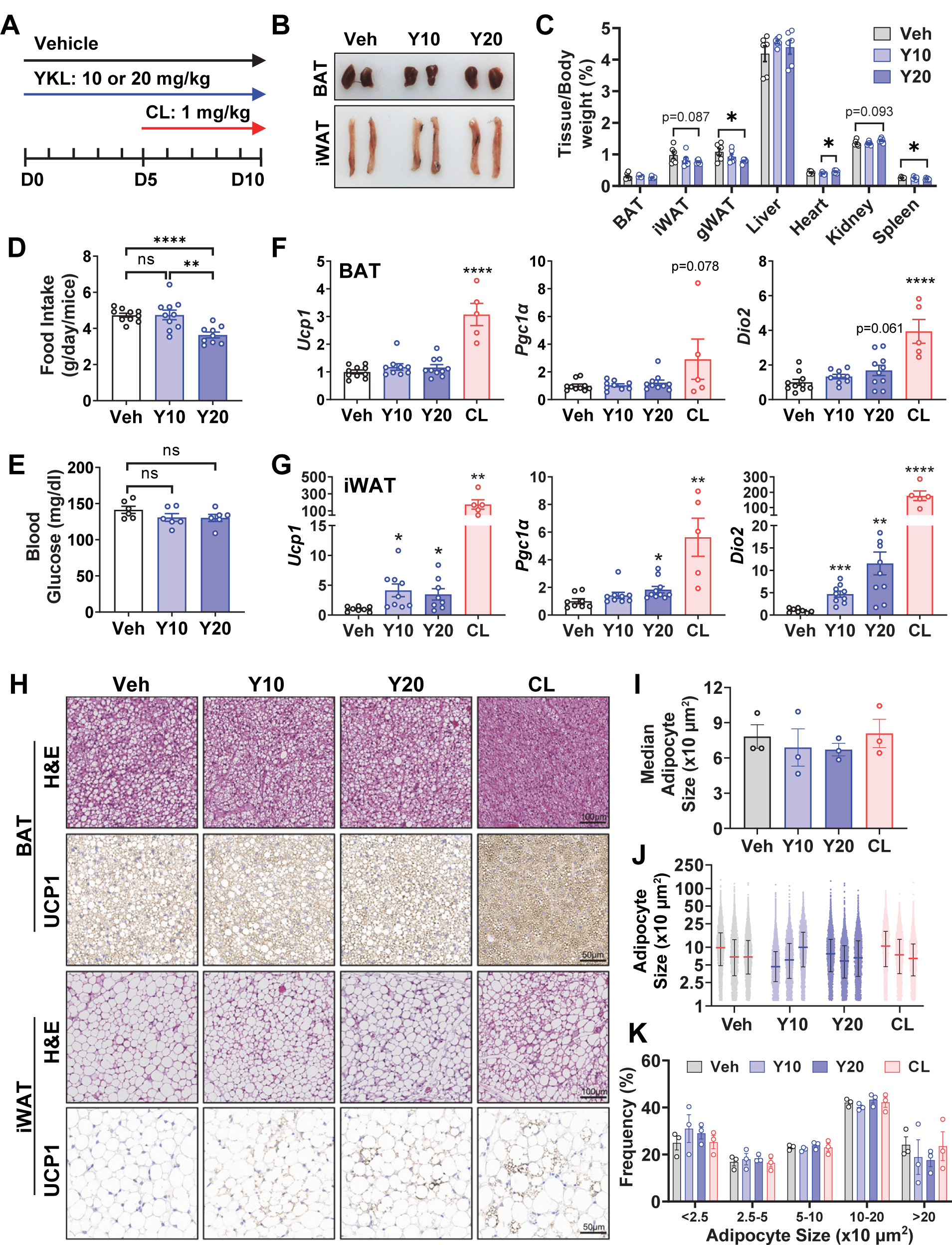
SIK inhibitor YKL promotes adipose tissue browning *in vivo*. (A) YKL was injected i.p. daily at 10 or 20 mg/kg bodyweight for 10 days. β_3_AR selective agonist CL was injected i.p. daily at 1 mg/kg bodyweight for 5 days as positive control. (B) Representative images of BAT and iWAT of control (Veh) or YKL treated mice (Y10, Y20). (C-E) Tissue to body weight ratio, daily food intake, and fasting blood glucose of control (Veh) and YKL treated mice (Y10, Y20). Student’s t-test, *p<0.05, **p<0.01, ***p<0.001, ns: no statistical difference. (F-G) QPCR analysis of *Ucp1*, *Pgc1α* and *Dio2* mRNA expression in BAT and iWAT of control (Veh), YKL (Y10, Y20), and CL-treated mice. Student’s t-test, *p<0.05, **p<0.01, ***p<0.001 vs vehicle control. (H) H&E and UCP1 immunohistochemistry staining of BAT and iWAT sections of control (Veh), YKL (Y10, Y20), or CL-treated mice. (I-K) Median adipocyte size, full range adipocyte size, and adipocyte size frequency in iWAT of control (Veh), YKL (Y10, Y20), or CL-treated mice.

The expression of genes representative of brown/beige adipocytes were assessed from tissues of these mice. The mRNA levels of adipose browning marker genes *Ucp1*, *Pgc1α* and *Dio2* (iodothyronine deiodinase 2) were increased in the subcutaneous iWAT, but not in the interscapular BAT of 10 mg and 20 mg treatment groups (**Figure 5F** and **5G**). Similarly, the adipose browning phenotype was further confirmed by the appearance of UCP1-positive adipocytes in the iWAT of 10 mg and 20 mg treatment group as indicated by immunohistochemical UCP1 staining (**Figure 5H**). However, the adipocyte size did not change in the iWAT of both 10 mg and 20 mg treatment group, as indicated by morphological segmentation analysis (**Figure 5I-5K**). Since the adipose tissue browning phenotypes in the YKL treatment groups occurred at room temperature without much additional β-adrenergic stimulation, it was not surprising that these phenotypes were not as robust as the CL treatment group (**Figure 5F-5H**). Nevertheless, these results demonstrated the potential of SIK inhibitor to improve adipose tissue browning *in vivo*. It will be informative to assess the whole-body metabolic effect when newer isoform selective SIK inhibitors are available in future.

## 4. DISCUSSION

In this study, we identified SIK3 as a novel substrate for mTORC1 in a SILAC study of βAR-specific signaling in brown adipocytes. Our study defines a role for SIK3 as a suppressor of the brown adipocyte thermogenic gene program. The regulatory PKA phosphorylation domain is essential for SIK3 inhibition to allow for the thermogenic gene expression and HDAC4 might be a downstream mediator of this process. Our study further demonstrated that pharmacological SIK inhibition promotes adipose tissue browning *in vivo*, even in the absence of additional βAR stimulation.

In line with previous reports [52; 53], our study suggested that SIKs might play redundant roles in the brown adipocyte thermogenic program. The shRNA knockdown of either *Sik1*, *Sik2*, or *Sik3* increased *Ucp1* expression in HIB-1B brown adipocytes. The pan-SIK inhibitors HG and YKL also induced *Ucp1* expression in brown adipocytes in the absence of additional βAR stimulation. Pharmacological SIK inhibition *in vivo* by YKL increased thermogenic gene expression and adipose browning *in vivo*. All three SIK isoforms are expressed in rodent and human adipocytes and adipose tissues, with the expression of SIK2 being most abundant [52; 54]. SIK2 has been previously linked to the regulation of insulin signaling, adipogenesis and lipogenesis in adipocytes [55–58]. In adipose tissue from obese or insulin-resistant individuals, SIK2 and SIK3 mRNAs have been reported to be down-regulated [54], but the functional consequences, as well as cell types affected, are still unclear. In mouse models, *Ucp1* expression is increased in iWAT of *Sik1/2* double knockout mice, but not in *Sik1* or *Sik2* single knockout mice [52]. Global *Sik3* deletion disturbs glucose and lipid metabolism in mice [59], however, these phenotypes could be also caused by defects in other metabolically relevant tissues where they are expressed, such as the liver. Further characterization of a genetic model with adipose-specific *Sik3* deletion will be necessary to define its role in regulating adipose tissue metabolism *in vivo*.

In addition, our study demonstrated that SIK3 lacking an internal PKA regulatory domain is resistant to PKA-mediated SIK inhibition, suggesting that phosphorylation by PKA is essential for SIK3 inhibition to drive the thermogenic gene expression after β-adrenergic stimulation. In this regard PKA inhibition of SIK3 might be a component of the overall effect of cAMP/PKA signaling to drive the program of browning and non-shivering thermogenesis. Our study further showed that SIK3 physically interacts with mTORC1 through RAPTOR, in line with a previous study demonstrating that SIK3 can interact with several components of the mTOR complex, including RAPTOR [60]. Although this latter study was done in the context of bone development, both studies suggest a connection between SIK3 and the mTOR complex, which might be applicable to other tissues or organs as well. Our SILAC study showed that S^884^ in SIK3 is phosphorylated after Iso stimulation in a mTORC1-dependent manner. The functional significance of this SIK3-mTORC1 connection needs to be further explored in the context of adipocyte thermogenesis and metabolism perhaps by taking more unbiased approaches to uncover facets of SIK3 regulation of beige/brown adipocytes that are not yet known. Our data showed that phosphorylation of S^884^ in SIK3 by mTORC1 alone does not change its function with respect to the regulation of thermogenic gene expression; it may function in synergy with other phosphorylation events to control SIK3 activity or its subcellular location.

As an established mechanism for the control of PGC1α coactivator activity, acetylation of the protein has been shown to be regulated by the histone acetyltransferase GCN5, the nicotinamide adenosine dinucleotide (NAD)-dependent deacetylase SIRT1, and N-α-acetyltransferase 10 (NAA10) [16–18]. Our results showed that SIK inhibition stimulates thermogenic gene expression through the action of HDAC4. We found that HDAC4 interacts with PGC1α after Iso stimulation and increase the deacetylation of PGC1α. Of note, another recent study also identified PGC1α as a new HDAC4 substrate in the control of skeletal muscle homeostasis [48]. This study, together with ours, suggest that HDAC4 might be a new regulator of PGC1α acetylation in metabolically active tissues such as the skeletal muscle and brown fat. The function of this HDAC4-PGC1α axis in the other relevant tissues and physiological contexts needs to be further explored in the future.

Several compounds have been reported as pan-SIK inhibitors, including HG [40] and YKL [42]. And isoform-selective SIK inhibitors have also been reported, such as the SIK2-selective inhibitor ARN-3236 [61; 62], and the SIK3-selective inhibitor PTB [43; 44]. Among these currently available inhibitors, the SIK inhibitor YKL has been shown *in vivo* to suppress acute myeloid leukemia progression [51], increase bone formation [29; 63], and modulate cytokine production [42]. Here our study showed that YKL stimulates thermogenic gene expression and promotes adipose tissue browning *in vivo*. While it is unclear whether SIK inhibition in the adipose tissue alone or in other cell type(s) or tissue(s) contribute to the adipose tissue phenotypes, and which SIK isoform(s) is primarily responsible for this effect on adipose browning, the result is nevertheless encouraging from a target therapy perspective. It supports a role for SIK inhibition in adipose tissue and represents a potential intervention strategy that could be translated to obesity and cardiometabolic disease in future studies. Parenthetically, newer inhibitors currently in development are more isoform selective and potent [64–66]. The newer SIK3-selective inhibitors will allow further mechanistic studies both *in vitro* and *in vivo* [66].

In summary, our data reveal that salt-inducible kinases function as a phosphorylation switch for β-adrenergic activation to drive the adipose thermogenic program downstream of mTORC1 activation. SIK3 deletion increases the thermogenic gene and mitochondrial protein expression in brown adipocytes. Pharmacological SIK inhibition by YKL promotes adipose tissue browning *in vivo*. Our study supports the notion that maneuvers targeting SIKs could be beneficial for obesity and related cardiometabolic disease.

## AUTHOR CONTRIBUTIONS

Conceptualization, FS, MK, and SC; Formal analysis, FS, CT, MK, and SC; Investigation, FS, FFS, DL, CT, HUP, JX, WZ; Resources, SC; Data curation, FS, CT; Writing – Original Draft, FS; Writing – Review & Editing, FS and SC; Visualization, FS, CT, MK, SC; Supervision, SC; Project administration, SC; Funding acquisition, SC; All authors approved the manuscript for publication.

## DATA AVAILABILITY

All data and reagents are available on request.

## ACKNOWLEDGEMENT

This work was supported by NIH R01 DK116625 (SC). FS is supported by the American Heart Association Career Development Award 23CDA1048341. We acknowledge the Vanderbilt Translational Pathology Shared Resource supported by NCI/NIH Cancer Center Support Grant 5P30CA68485 and the Shared Instrumentation Grant S10OD023475. The Agilent Seahorse Extracellular Flux Analyzer is housed and managed within the Vanderbilt High-Throughput Screening Core Facility, an institutionally supported core, and was funded by NIH Shared Instrumentation Grant 1S10OD018015.

### APPENDIX A. SUPPLEMENTARY DATA

Supplementary Table 1: Plasmid List.

Supplementary Table 2: QPCR Primers.

## Abbreviations

BAT: brown adipose tissue
WAT: white adipose tissue
SIK: salt-inducible kinase
SILAC: Stable Isotope Labeling by/with Amino acids in Cell culture
βAR: β-adrenergic receptor
cAMP: cyclic adenosine monophosphate
PKA: protein kinase A
mTORC1: mechanistic target of rapamycin complex 1
HDAC: histone deacetylase
CREB: cAMP response element-binding protein
CRTC: CREB regulated transcription coactivator

