## Supplemental Data for "Salt-inducible kinase inhibition promotes the adipocyte thermogenic program and adipose tissue browning"

**Supplementary Table 1: Plasmid List.**

| **Plasmid** | **Description** | **Application** |
| --- | --- | --- |
| SIK3-Flag | pCMV6-Neo-3xFlag-SIK3 | Co-expression, subcloning |
| SIK3-Flag-ΔPKA | pCMV6-Neo-3xFlag-SIK3-ΔPKA | Subcloning |
| SIK3-Flag-ΔKD/PKA | pCMV6-Neo-3xFlag-SIK3-ΔKD/PKA | Subcloning |
| SIK3-Flag-S884A | pCMV6-Neo-3xFlag-SIK3-S883A | Subcloning |
| SIK3-Flag-S884D | pCMV6-Neo-3xFlag-SIK3-S883D | Subcloning |
| Lenti-SIK3-Flag | pLenti-Puro-3xFlag-SIK3 | Over-expression |
| Lenti-SIK3-Flag-ΔPKA | pLenti-Puro-3xFlag-SIK3-ΔPKA | Over-expression |
| Lenti-SIK3-Flag-ΔKD/PKA | pLenti-Puro-3xFlag-SIK3-ΔKD/PKA | Over-expression |
| Lenti-SIK3-Flag-S884A | pLenti-Puro-3xFlag-SIK3-S883A | Over-expression |
| Lenti-SIK3-Flag-S884D | pLenti-Puro-3xFlag-SIK3-S883D | Over-expression |
| shRNA-mSik1 | pLKO.1-Puro-mSik1 | Knock-down |
| shRNA-mSik2 | pLKO.1-Puro-mSik2 | Knock-down |
| shRNA-mSik3 | pLKO.1-Puro-mSik3 | Knock-down |
| shRNA-mHdac4 | pLKO.1-Puro-mHdac4 | Knock-down |
| shRNA-mHdac5 | pLKO.1-Puro-mHdac5 | Knock-down |
| shRNA-mHdac7 | pLKO.1-Puro-mHdac7 | Knock-down |
| shRNA-mPgc1α | pLKO.1-Puro-mPgc1α | Knock-down |
| NAA10-Myc | pCMV6-Neo-Myc-NAA10 | Co-expression |

**Supplementary Table 2: QPCR Primers.**

| **QPCR** | **Fwd (5’-3’)** | **Rev (5’-3’)** |
| --- | --- | --- |
| 36B4 | GATGCCCAGGGAAGACAG | ACAATGAAGCATTTTGGATAATCA |
| Ucp1 | GGCCTCTACGACTCAGTCCA | TAAGCCGGCTGAGATCTTGT |
| Pgc1α | CGGAAATCATATCCAACCAG | TGAGAACCGCTAGCAAGTTTG |
| Sik3 | CCAGAGCTCTTCGAAGGGAA | CAGTGTGCTCCCATCAAACG |
| Dio2 | gcctctcaggacagaagtgc | ctgccaaagttcaacaccag |
| Hdac4 | gcaagatcctcattgtagactgg | gaacattggggtcattgtagaag |
| Hdac5 | gcactggagggtgacacaa | acgttcacccgtcaccag |
| Hdac7 | gcagcccttgagagaacagt | tgtccaagggctcaagagtt |
| Sik1 | tgcaggctgagatagactgtg | gctgatagctgtgtccagca |
| Sik2 | tccaagacctttcgagcagt | gcagacaggctggagacac |
